# Loss of growth differentiation factor 9 causes an arrest of early folliculogenesis in zebrafish – a novel insight into its action mechanism

**DOI:** 10.1101/2022.07.01.498398

**Authors:** Weiting Chen, Yue Zhai, Bo Zhu, Kun Wu, Yuqin Fan, Xianqing Zhou, Lin Liu, Wei Ge

## Abstract

Growth differentiation factor 9 (GDF9) is the best characterized growth factor released by the oocyte; however, most information about GDF9 functions comes from studies in the mouse model. In this study, we created a mutant for Gdf9 gene (*gdf9-/-*) in zebrafish using TALEN approach. The loss of Gdf9 caused a complete arrest of follicle development at primary growth (PG) stage. These follicles eventually degenerated, and all mutant females gradually changed to males through sex reversal, which could be prevented by mutation of the male-promoting gene *dmrt1*. Interestingly, the phenotypes of *gdf9-/-* could be rescued by mutation of inhibin α (*inha*-/-) but not estradiol, suggesting a potential role for the activin-inhibin system in Gdf9 actions. In *gdf9* null follicles, activin βAa (*inhbaa*) expression decreased dramatically; however, its expression rebounded in the double mutant (*gdf9*-/-*;inha-/-*). These results clearly indicate that although endocrine hormones such as follicle-stimulating hormone (FSH) are important for folliculogenesis, the activation of PG follicles requires intrinsic factors from the oocyte, such as Gdf9, which in turn works on the neighboring follicle cells to trigger follicle activation, probably via activins.

## Introduction

Follicles, each consisting of an oocyte and surrounding somatic follicle cells, are the basic structural and functional units of the ovary. The growth and maturation of follicles or folliculogenesis is primarily controlled by gonadotropins (follicle-stimulating hormone, FSH; and luteinizing hormone, LH) from the pituitary in both mammals and fish [1–4]. However, evidence has accumulated in various species that local ovarian factors also play critical roles in controlling folliculogenesis in paracrine and/or autocrine manners [5–9].

In the past two decades, there has been substantial progress in understanding the mechanisms underlying folliculogenesis, especially its early stage [10–12]. Since this stage of follicle development is considered less dependent on gonadotropins, most studies have focused on intraovarian growth factors that work locally through paracrine and/or autocrine pathways [10, 11, 13, 14]. Among numerous local factors characterized, those released by the oocyte have caught much attention in research, particularly growth differentiation factor 9 (GDF9) and bone morphogenetic protein 15 (BMP15) [15–18]. Malfunction of these factors has been implicated in some human reproductive disorders. For example, some POCS or POF cases in humans have been associated to mutations or expression abnormalities of GDF9 [19–21] or BMP15 [20, 22].

As a TGF-β family member, GDF9 was first discovered in 1993 as an ovarian growth factor [23]. Subsequent studies demonstrated that the expression of GDF9 mRNA and protein was restricted exclusively to the oocyte in several mammalian species including mice and humans [25]. As a novel oocyte-specific factor, GDF9 is now recognized as a critical player to orchestrate folliculogenesis by signaling the surrounding follicle cells, namely the granulosa and theca cells [14, 26, 27]. The essential role of GDF9 in folliculogenesis was demonstrated in the GDF9 knockout mice, whose follicles could develop normally to the stage of primary follicles but not beyond, causing female infertility [28, 29]. Studies using recombinant GDF9 showed that GDF9 stimulated initial follicle recruitment in the rat ovary [30] and promoted progression of human primordial follicles to the secondary stage *in vitro* [31]. Both the granulosa and theca cells surrounding the oocyte are the targets of GDF9 action, and its effects vary during folliculogenesis depending on the stage of follicle development [27]. GDF9 promoted granulosa cell proliferation and FSH-induced cumulus expansion *in vitro*, and increased inhibin and progesterone secretion from these cells [30, 32–35]. GDF9 also stimulated the expression of a variety of genes in the granulosa cells including Kit ligand (KL) [36], cyclooxygenase 2 and steroidogenic acute regulator (StAR) protein [33]. In the theca cells, GDF9 seemed essential for the expression of theca cell-specific genes such as CYP17 and LH receptor [37, 38]. In cultured human granulosa-lutein cells, GDF9 alone showed weak stimulatory effects on expression of activin βA and βB subunits; but it dramatically increased the activin-induced βA and βB expression [39]. On the other hand, GDF9 inhibited FSH-induced steroidogenesis and LH receptor expression in the granulosa cells [30]. The physiological importance of GDF9 has also been documented by *in vivo* studies. In humans, a mutation in *GDF9* caused diminished ovarian reserve in young women [40]. In sheep, a missense *GDF9* mutation (V371M) was strongly associated with litter size [41].

Unlike other members of the TGFβ superfamily with well-documented serine/threonine kinase receptors (type I and II) and intracellular signaling pathways involving specific Smads, the signaling mechanism of GDF9 still remains elusive. Some studies have implicated bone morphogenetic protein (BMP) type II receptor (BMPR-II) in GDF9 signaling [42, 43], which recruits and activates a type I receptor ALK5 (activin receptor-like kinase 5) [44]. The activation of ALK5 in turns induces phosphorylation of Smad2 and Smad3 for intracellular signaling, which can be blocked by inhibitory Smad7, but not Smad6 [32, 39, 44, 45], a mechanism shared by TGFβ and activin.

Compared with studies in mammals especially mice, the information about GDF9 in other species remains limited. Our laboratory was the first to isolate *gdf9* gene in teleosts in 2007 and provided evidence for a potential role of this factor in controlling early follicle development in the zebrafish. As in mammals, zebrafish *gdf9* is also expressed exclusively in the oocyte. During zebrafish gonadal differentiation, which we divide into three stages: pre-differentiation (15-25 dpf, days post-fertilization), differentiation (25-35 dpf) and post-differentiation (35-45 dpf) [46], *gdf9* expression starts to increase as early as 16 dpf before morphological differentiation starts, suggesting a role for Gdf9 in ovarian formation [47]. In adult zebrafish, the expression of *gdf9* mRNA in females was the highest in the primary growth (PG) follicles, but decreased significantly when the follicles entered the secondary growth (SG) phase, suggesting a role for Gdf9 in controlling PG-SG transition or follicle activation [48]. Our recent study demonstrated that recombinant zebrafish Gdf9 could act on cultured follicle cells, stimulating Smad2 phosphorylation and expression of activin subunits (*inhbaa* and *inhbb*) but suppressing the expression of anti-Müllerian hormone (*amh*) [47]. A study using recombinant mouse GDF9 showed that GDF9 might regulate tight junction protein expression during zebrafish folliculogenesis [49], and immunization with synthetic zebrafish GDF9 peptide caused abnormal oocyte development [50]. In Japanese flounder, overexpression of *gdf9* in an ovarian cell line stimulated expression of most steroidogenic genes [51]. Despite these studies, the functional importance of GDF9 in fish has remained largely unknown due to the lack of genetic approaches such as gene knockout as in the mouse model.

In recent years, a new generation of gene editing technologies, in particular TALEN and CRISPR/Cas9, has revolutionized research in biological fields [52], and these technologies are now easily available in fish, in particular the model species such as zebrafish [53]. To test our previous hypothesis that GDF9 plays an important role in zebrafish folliculogenesis especially at early stage, we undertook this study to knock out *gdf9* gene in zebrafish using TALEN method. Our data strongly demonstrated essential roles for GDF9 (Gdf9/*gdf9*) in early follicle development, especially the activation of follicles to enter the secondary growth.

## Results

### Generation of gdf9 knockout zebrafish by TALEN

We generated *gdf9* mutant fish (*gdf9-/-*) by transcription activator-like effector nucleases (TALEN) method. The TALEN target site of *gdf9* is located in the first exon downstream of the ATG start codon. A mutant fish with 4-bp deletion in the *gdf9* gene was generated (*gdf9^-4/-4^*) (ZFIN line number: umo18), which resulted in a frameshifting mutation that produced a truncated protein with 33 amino acids of the Gdf9 precursor. The mutation was also confirmed at mRNA level in the ovary by RT-PCR using a mutant-specific primer with the 3’-end over the deleted sequence, which would generate a positive product in wild type (WT), but not the mutant fish (Fig.1A).

**Fig. 1.**
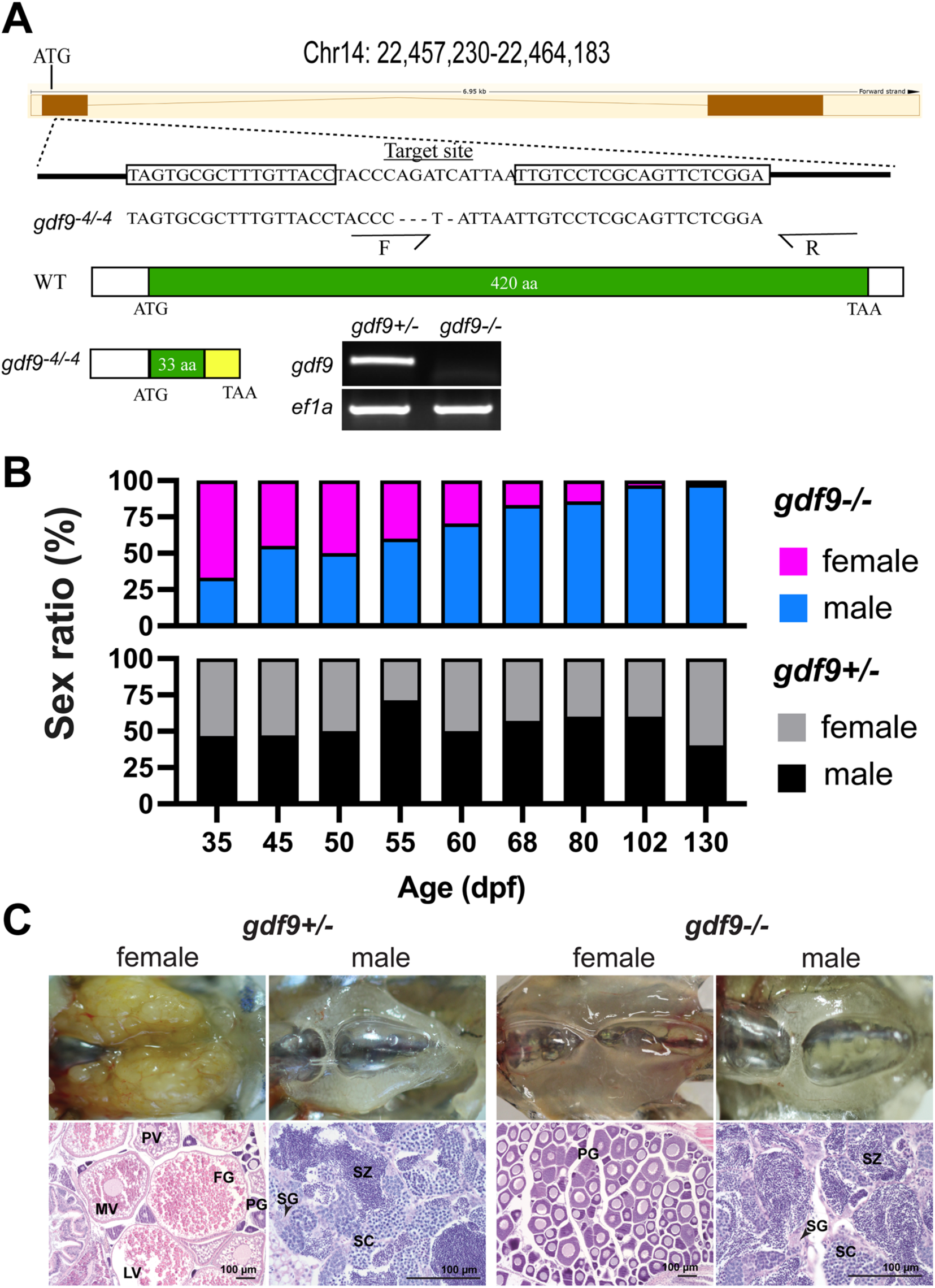
Disruption of *gdf9* gene in zebrafish and evidence for its roles in sex differentiation and gonadal development. (A) Generation of *gdf9* mutant by TALEN. The target site on chromosome 14 for left and right TALE proteins are boxed. A mutant with 4-bp deletion was obtained with a premature stop codon (TAA), which generated a truncated protein with 33 amino acids of the Gdf9 precursor. RT-PCR analysis on mRNA from the ovary using a mutant-specific primer (F) showed a positive signal in the control ovary (*gdf9*+/-), but not in the mutant (*gdf9*-/-). (B) Sex ratios during and after gonadal differentiation in *gdf9* mutant fish. The sex ratios in the post-differentiation period from 35 to 55 dpf remained relatively constant and comparable to those in the control fish. However, the ratio changed gradually but significantly afterwards towards all males. (C) The ovary and testis in the control (*gdf9+/-*) and mutant (*gdf9-/-*) at 90 dpf. The control females had reached sexual maturity with all stages of follicles present in the ovary (PG, primary growth; PV, pre-vitellogenic; MV, mid-vitellogenic; LV, late vitellogenic; and FG, full-grown) while the mutant ovary contained PG follicles only. The testis development and spermatogenesis showed no difference between mutant and control males. SG, spermatogonia; SC, spermatocytes; SZ, spermatozoa.

### Role of Gdf9 in gonadal differentiation

Gdf9 is an oocyte-specific growth factor in both mammals and fish [24, 48] and its expression in zebrafish increases significantly when the gonads differentiate towards ovaries during sex differentiation together with ovarian aromatase (*cyp19a1a*) [47]. This had led us to hypothesize that Gdf9 might play a critical role in driving ovarian differentiation in zebrafish. To test this hypothesis, we examined sex ratio and its changes during and after gonadal differentiation from 35 to 130 dpf. To our surprise, the lack of Gdf9 did not have any impact on sex differentiation. The sex ratios of the mutant (*gdf9-/-*) were normal and comparable to those in the age-matched control fish (*gdf9+/-*) (∼50%) during the post-differentiation period from 35 to 55 dpf. However, the female ratio declined progressively afterwards, and nearly all fish examined were males at 130 dpf except a few still containing some oocytes in the gonads (∼3%), suggesting sex reversal from females to males (Fig. 1B).

### Arrest of follicle development at PG stage in gdf9 mutant females

To find out gonadal status in *gdf9* mutant (*gdf9-/-*), we first examined mature males and females at 90 dpf using *gdf9+/-* as the control (we did not observe any difference between *gdf9+/+* and *gdf9+/-*). The control fish already reached maturity with all stages of follicles present in the ovary and full scale of spermatogenesis in the testis. The spermatogenesis in the mutant testis seemed normal compared to the control with abundant mature spermatozoa in the tubular lumen. However, the follicles in the mutant ovary were completely arrested at the PG stage without any signs of transition to the SG phase, which begins with the pre-vitellogenic (PV) stage characterized with formation of cortical alveoli (vesicles) (Fig. 1C).

To follow the developmental process of gonads after sex differentiation, we performed a systematic analysis on both ovary and testis by histology from gonadal differentiation to sexual maturation (35-80 dpf). At 35 dpf, the gonads had completed differentiation with ovary and testis well formed in both control (*gdf9+/-*) and mutant (*gdf9-/-*) fish. The ovaries contained PG follicles (stage I) and the testes were at pre-luminal stage (PL; stage I) with signs of meiosis according to our recent categorization of spermatogenic stages [54]. At 45 dpf, the leading follicles in the control females began accumulating cortical alveoli in the oocytes (PV; stage II), a morphological marker for follicle activation and transition from PG to SG phase (PG-PV transition) [55]. These follicles continued to grow through vitellogenic growth (from early vitellogenic to full-grown, EV-FG; stage III) to become mature from 55 to 80 dpf. In contrast, the follicles in the mutant ovary remained arrested at PG throughout the period without any signs of follicle activation such as formation of cortical alveoli in the oocytes. At 80 dpf, the control females had reached sexual maturity with all stages of follicles present in the ovary. However, the mutant ovaries were undergoing degeneration; the follicles were well separated and much part of the ovary including interfollicular spaces was occupied by infiltrating stromal tissues, a sign of masculinization and sex reversal (Fig. 2A). What should be noted is that although most mutant females were arrested at the PG stage, the follicles in some individuals were able to enter very early PV stage with formation of rudiment cortical alveoli; however, this was not common (Fig. 2B).

**Fig. 2.**
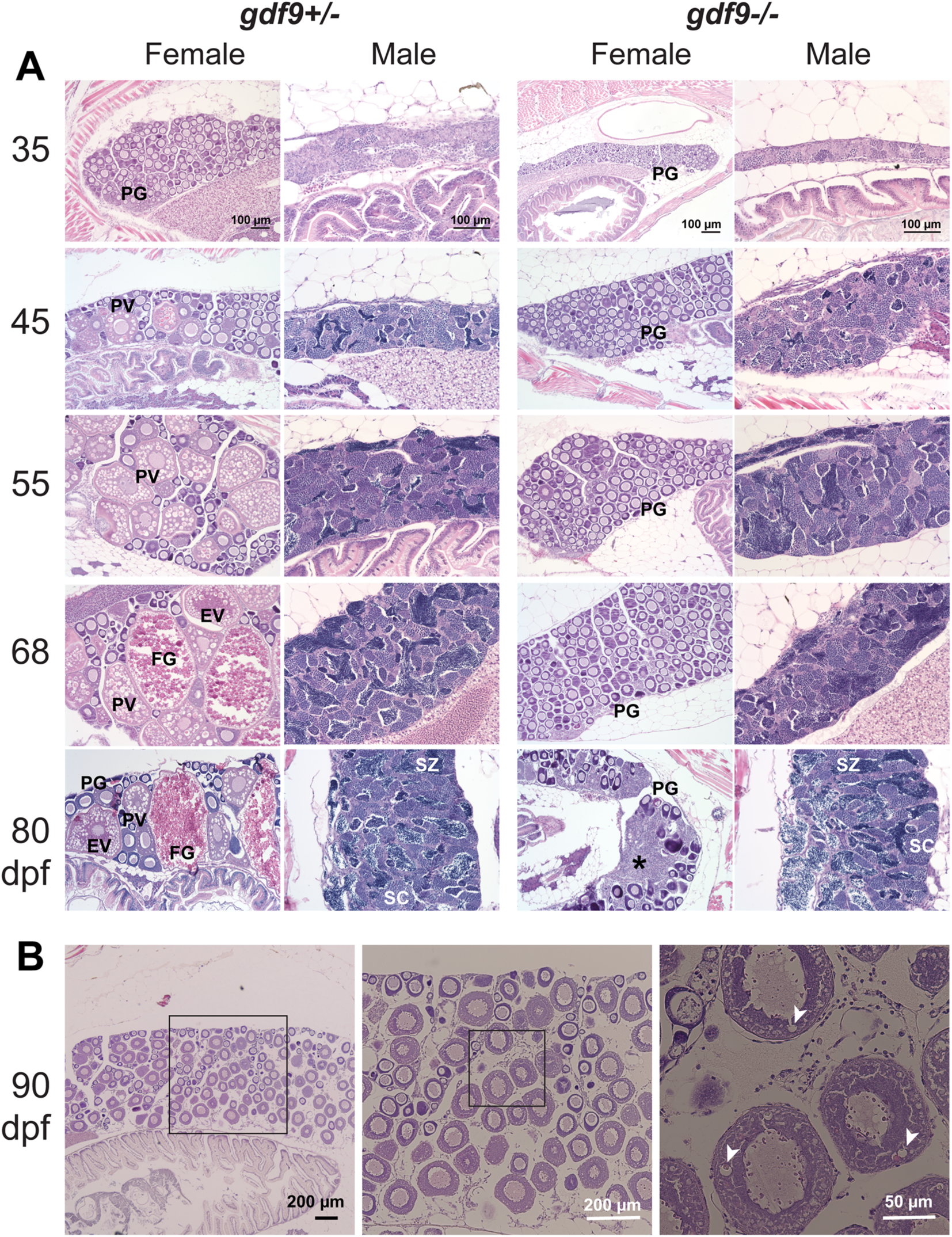
Gonadal development from sex differentiation to maturity in *gdf9* mutant. (A) Temporal development of the ovary and testis in the control (*gdf9+/-*) and mutant (*gdf9-/-*) from sex differentiation (35 dpf) to puberty onset (45 dpf) and sex maturity (80 dpf). The mutant females started to differ from the control at 45 dpf when the first cohort of PV follicles with cortical alveoli emerged in the control ovary, marking the onset of puberty. No PV follicles were observed in the mutant ovary, and the follicles remained arrested at PG stage during the entire examination period. At 80 dpf, the mutant females were undergoing sex reversal to males with remnant follicles and infiltrating stromal and testicular tissues (asterisk). The testis and spermatogenesis were normal in the mutant males compared to the control. (B) Very few *gdf9* mutant females could slightly cross the PG-PV transition and their follicles showed rudiment cortical alveoli (arrowhead). PG, primary growth; PV, pre-vitellogenic; EV, early vitellogenic; FG, full-grown; SC, spermatocytes; SZ, spermatozoa.

As for males, although *gdf9* expression could also be detected in adult testis [48], we did not see any abnormalities in testis development and spermatogenesis in mutant males (*gdf9-/-*) at any time points of examination compared to age-matched control males. At 45 dpf, the testes in both control and mutant started to produce small amount of mature spermatozoa in the tubular lumen (mid-luminal stage, ML; stage III), which we defined as the marker for puberty onset in males [54]. The post-pubertal spermatogenesis in the mutant males remained normal throughout the examination period compared to the control (Fig. 2A).

### Sex reversal of gdf9 mutant from female to male

The change of sex ratio after post-differentiation period suggested that the female mutant (*gdf9-/-*) might undergo sex reversal from females to males after being arrested at PG stage for some time. This was confirmed by histological analysis. The mutant ovary started to degenerate after a long time of follicle arrest at PG stage. While the ovarian tissues/follicles were degenerating and receding, the stromal tissues containing gonial or early meiotic germ cells (spermatogonia) gradually infiltrated the interfollicular spaces followed by emergence of testicular tissues with active spermatogenesis (Fig. 3). The process of sex reversal began around 45 dpf or earlier, and it took about 2-3 months for all female mutants to change to males (Fig. 1B and 2A). The sex-reversed males were functional with normal spermatogenesis and they could spawn with WT females to produce normal offspring.

**Fig. 3.**
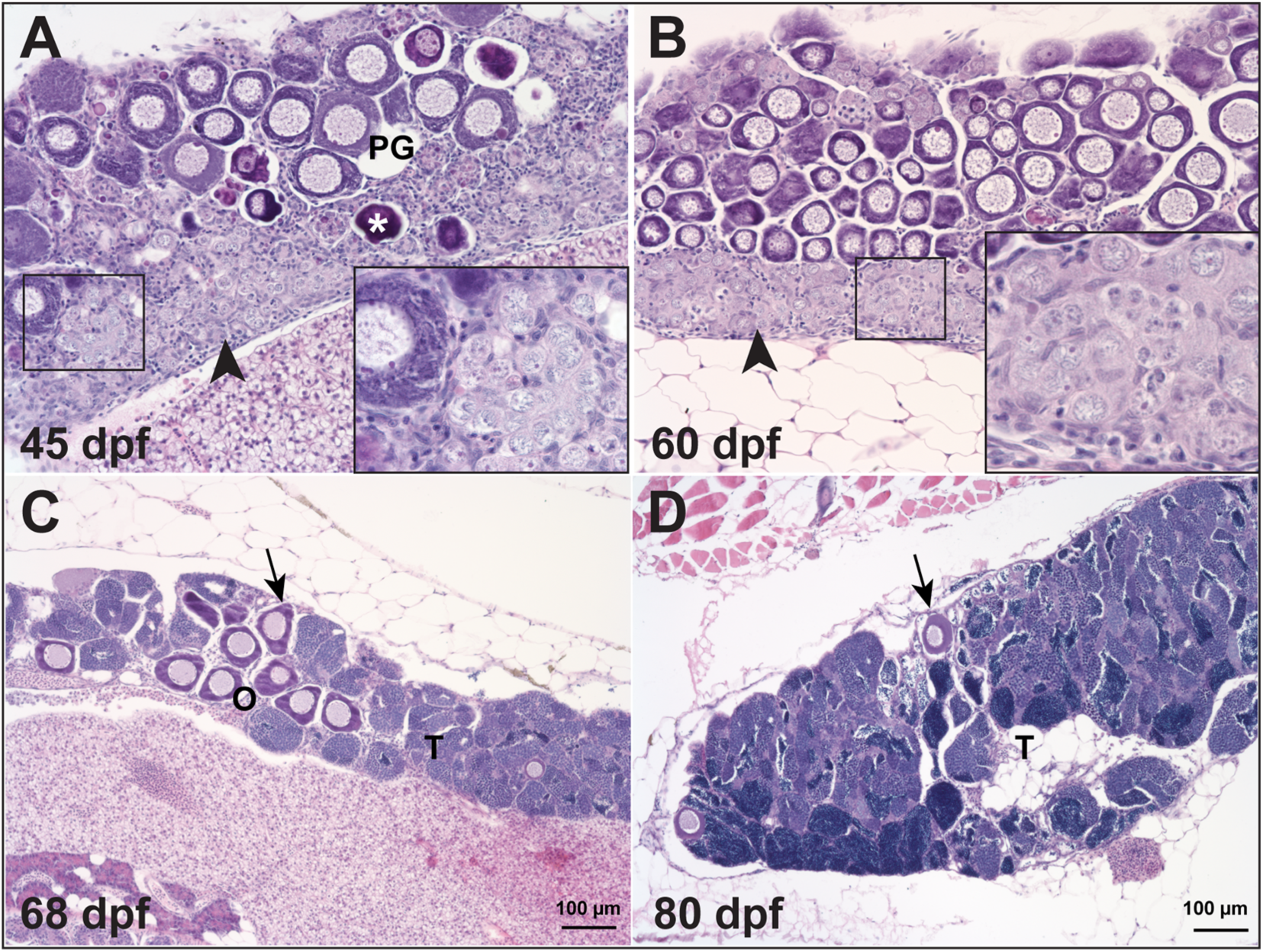
Temporal process of sex reversal from females to males. Sex reversal in *gdf9* mutant started around 45 dpf when the follicles were undergoing degeneration (asterisk) while stromal tissue with early spermatogenic cells (inset) was infiltrating the ovarian tissue (O), which normally started from the ventral side of the ovary (arrowhead). The number of PG follicles (arrow) decreased progressively during the sampling period (45-80 dpf) with increasing amount of testicular tissue (T).

Recently we demonstrated that the loss of ovarian aromatase (*cyp19a1a*-/-) also resulted in all-male phenotype, which was due to the failure of ovarian differentiation and sex reversal from juvenile ovary to testis [56]. Surprisingly, double mutation of *cyp19a1a* and male-promoting gene *dmrt1* (*cyp19a1a-/-;dmrt1-/-*) prevented sex reversal from juvenile females to males in *cyp19a1a* mutant and therefore rescued the all-male phenotype of *cyp19a1a*-/-, resulting in normal sex ratio and normal ovarian formation [57]. To see if the loss of *dmrt1* has any effect on the sex reversal in *gdf9* mutant, we created a double mutant (*gdf9-/-;dmrt1-/-*) and examined its gonadal development up to 190 dpf. Interestingly, mutation of *dmrt1* (*dmrt1-/-*) (ZFIN line number: umo15) also prevented sex reversal in *gdf9* mutant (*gdf9-/-*) with the double mutant (*gdf9-/-;dmrt1-/-*) showing a female-biased sex ratio and the ovaries containing PG follicles only without any signs of sex reversal such as infiltrating stromal tissues (Fig. 4).

**Fig. 4.**
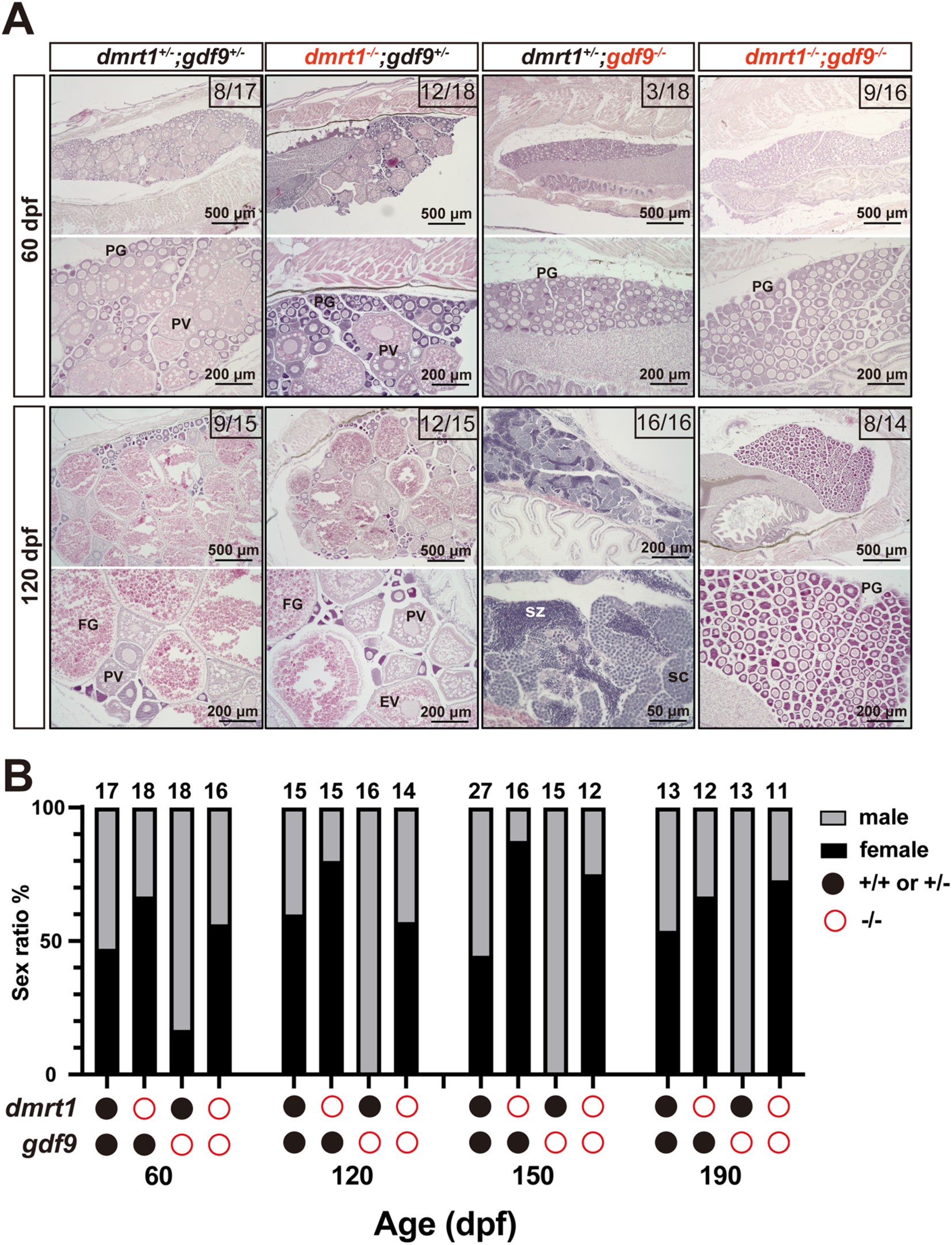
Prevention of sex reversal of *gdf9* mutant by *dmrt1* mutation. (A) Gonadal histology at 60 and 120 dpf. Ovary could form normally in the control (*dmrt1*+/-*;gdf9*+/-), single mutants (*dmrt1*-/- and *gdf9*-/-) and double mutant (*dmrt1*-/-*;gdf9*-/-) as seen at 60 dpf. The *gdf9*-/- single mutant became all males at 120 dpf (16/16 fish examined in total) whereas most individuals of the double mutant were females (8/14 examined), similar to that in the control (9/15). (B) Change of sex ratio during 60-190 dpf. The *gdf9*-/- fish showed a male-biased sex ratio at 60 dpf, and it became all-male from 120 dpf due to sex reversal. Double mutation with *dmrt1* rescued the sex ratio to the control level with even higher female ratios at 150 and 190 dpf. PG, primary growth; PV, pre-vitellogenic; FG, full-grown; SC, spermatocytes; SZ, spermatozoa.

### Expression analysis of key regulatory factors in the follicles of gdf9 mutant

To investigate the mechanism underlying Gdf9 actions, we examined the expression of several key ovarian factors at 45 dpf when some follicles were entering the PV stage in the control fish. We isolated the PG and early PV follicles from the control fish (*gdf9+/-*) and PG follicles from *gdf9-/-* fish (no PV follicles in the mutant) to analyze expression of genes by real-time qPCR, including gonadotropin receptors (*fshr* and *lhcgr*), ovarian aromatase (*cyp19a1a*), epidermal growth factor receptor (*egfra*) and members of the activin-inhibin family (inhibin α subunit *inha*, activin β subunits *inhbaa*, *inhbab* and *inhbb*). Most of these genes showed increased expression during the PG-PV transition in the control fish (*fshr*, *cyp19a1a*, *egfra*, *inha*, *inhbaa* and *inhbab*) except *lhcgr* and *inhbb,* which agreed well with our previous studies [58–62]. However, their expression showed little change in mutant PG follicles compared to stage-matched PG follicles from the control except activin βAa (*inhbaa*). Among all the genes examined, *inhbaa* was the only one that decreased expression dramatically in the mutant PG follicles (*gdf9-/-*) compared to the PG follicles from the control (*gdf9+/-*) (Fig. 5).

**Fig. 5.**
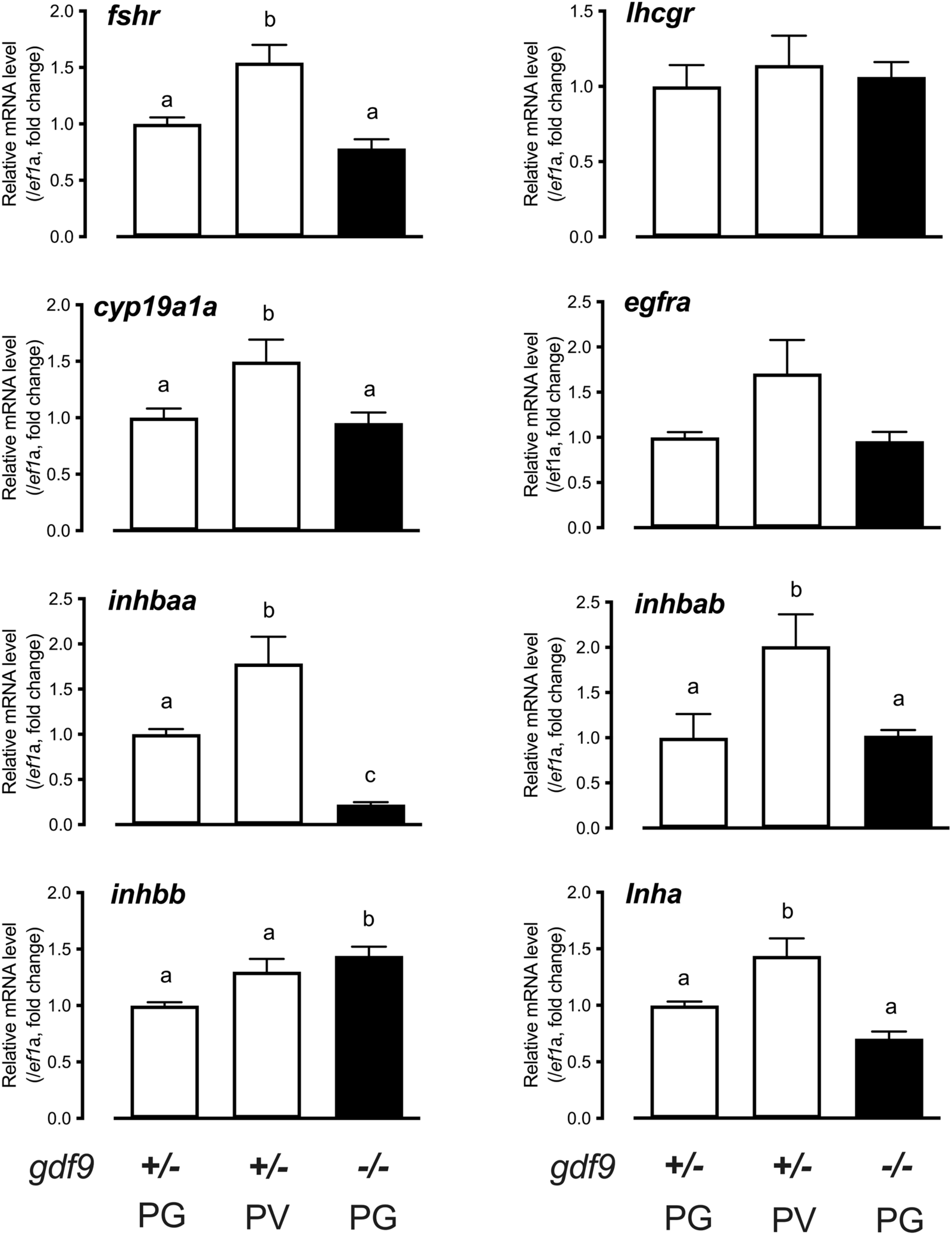
Gene expression in the control and mutant follicles at 45 dpf. Follicles were isolated from the control (*gdf9+/-*) (PG and PV) and mutant (*gdf9-/-*) (PG only) for total RNA extraction and real-time qPCR analysis. The data were normalized to the house-keeping gene *ef1a* and then expressed as the fold change over the control PG. Different letters indicate statistical significance (P < 0.05). *fshr,* FSH receptor; *lhcgr*, LH receptor; *cyp19a1a*, ovarian aromatase; *egfra*, EGF receptor a; *inhbaa*, activin βAa; *inhbab*, activin βAb; *inhbb*, activin βB; *inha*, inhibin α.

### Effect of estrogen exposure on follicle development in gdf9 mutant females

During PG-PV transition, the expression of *cyp19a1a* increased significantly [62]. To test if the arrest of follicles at PG stage in *gdf9* null females was due to the lack of sufficient estrogens, we treated the juvenile mutant fish (*gdf9-/-*) with estradiol (E2) either via water-borne exposure (10 nM) for 20 days (40-60 dpf) or oral administration (feeding) (20 µg/g diet) for 17 days (45 to 62 dpf). E2 administration via water-borne exposure induced all-female sex ratio (100% females), in contrast to the male-biased sex ratio in the control (∼70% males) (Fig. 6A). Furthermore, E2 also significantly increased the vitellogenin levels in both serum and liver of the mutants (*gdf9*-/-), especially the serum level (Fig. 6B). However, histological examination showed that E2 had little effect on follicle blockade in mutant females except that some follicles showed rudiment signs of cortical alveoli (Fig. 6C), the same as what we occasionally observed in some control fish (*gdf9+/-*) (Fig. 2B). Similar to water-borne exposure, oral administration of E2 by feeding had no effect on follicle activation either. All follicles remained arrested at the PG stage without any signs of transition to PV stage (Fig. 6D).

**Fig. 6.**
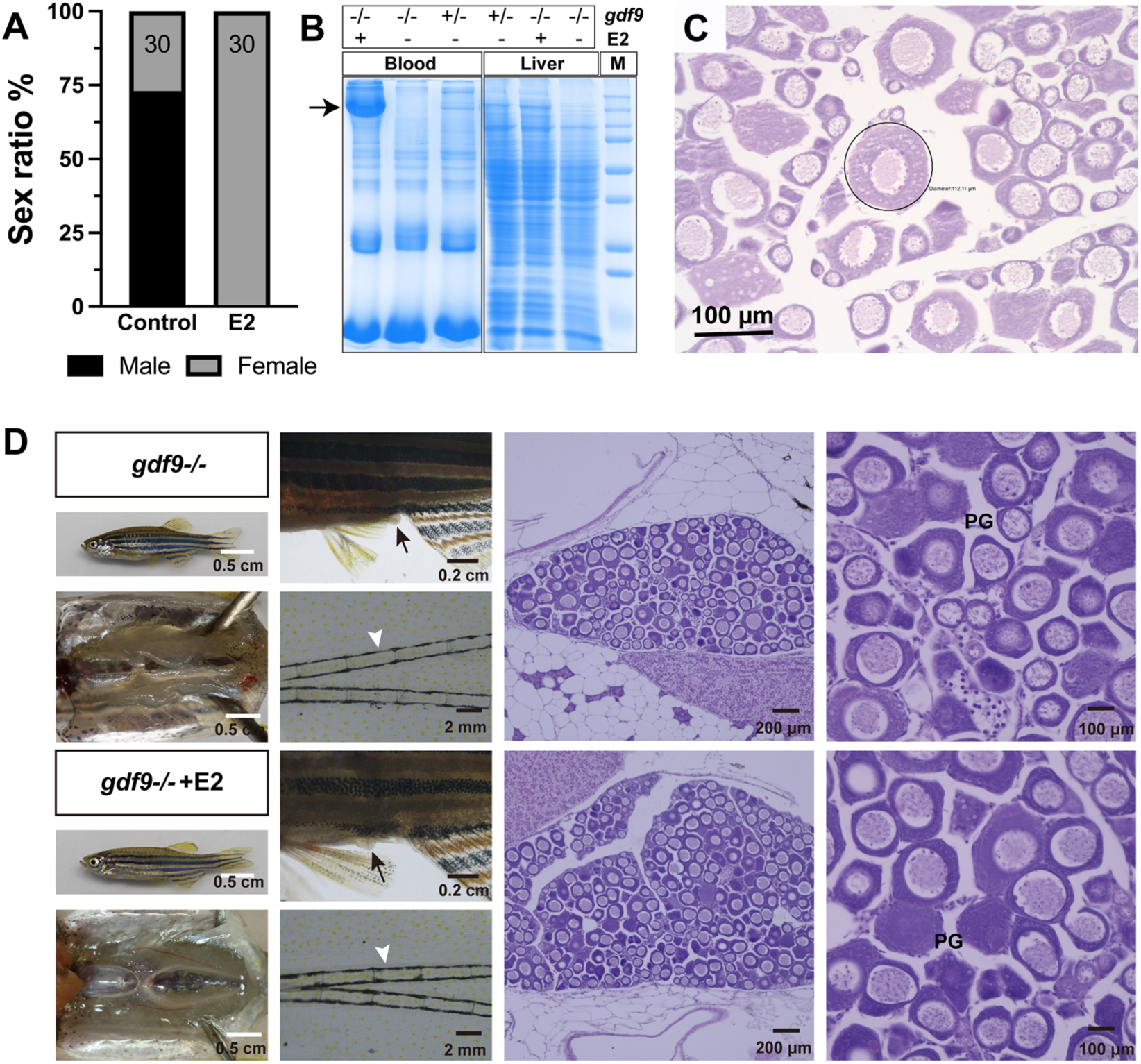
Estrogen treatment of *gdf9* mutant. (A-C) Treatment with estradiol (E2) by water-borne exposure. The mutant fish (30/group) were treated from 40 to 60 dpf with E2 (10 nM). E2 treatment resulted in all-female phenotype in contrast to untreated group which showed a male-biased sex ratio (A). Electrophoresis of the sera (1 µl/lane) from treated mutant females showed mass production of vitellogenin protein (arrow); in contrast, there was no vitellogenin in the sera of untreated mutant fish (-/-). The control fish (+/-) without E2 treatment showed a weak band at the location of vitellogenin. A similar band could also be seen in the liver extract (10 µg/lane) in response to E2 treatment (B). Histology of the ovary from E2-treated fish showed arrest of follicles at PG stage with a few follicles (circled) showing rudiment cortical alveoli in some individuals (C). (D) Treatment with E2 by oral administration (feeding). The mutant fish were fed with E2-containing diet (20 µg/g) for 17 days (45-62 dpf) followed by examination for secondary sex characteristics (genital papilla in females, arrow; breeding tubercles in males, arrowhead) and gonadal histology. The treated group showed higher female ratio (44% females, 8/18) compared to the control group (21% females, 3/14). Females from both groups showed normal genital papilla with no breeding tubercles and arrested PG follicles in the ovary. PG, primary growth.

### Rescue of gdf9 mutant phenotypes by inha deficiency

The dramatic decrease of activin βAa (*inhbaa*) in *gdf9* mutant (*gdf9-/-*) led us to hypothesize that the activin-inhibin system could be part of the underlying mechanism for Gdf9 actions in zebrafish follicle development. To test this hypothesis, we made a double mutant fish (*gdf9-/-;inha-/-*) by crossing the *gdf9* mutant with an *inha* mutant we reported recently (umo19) [63]. Surprisingly, the loss of inhibin, a natural activin antagonist, could significantly rescue the phenotypes of *gdf9* mutant. Without inhibin (*inha-/-*), the follicles arrested in *gdf9* mutant (*gdf9-/-*) could undergo the PG-PV transition to enter the fast SG phase. As the marker for puberty onset, cortical alveoli could form normally in the oocytes (PV stage) followed by accumulatio of yolk granules (vitellogenic stage). The double mutant (*gdf9-/-;inha-/-*) also phenocopied *inha* single mutant (*inha-/-*) in that the yolk mass in the oocytes had a sharp boundary compared to that in the control oocytes (Fig. 7) [63].

**Fig. 7.**
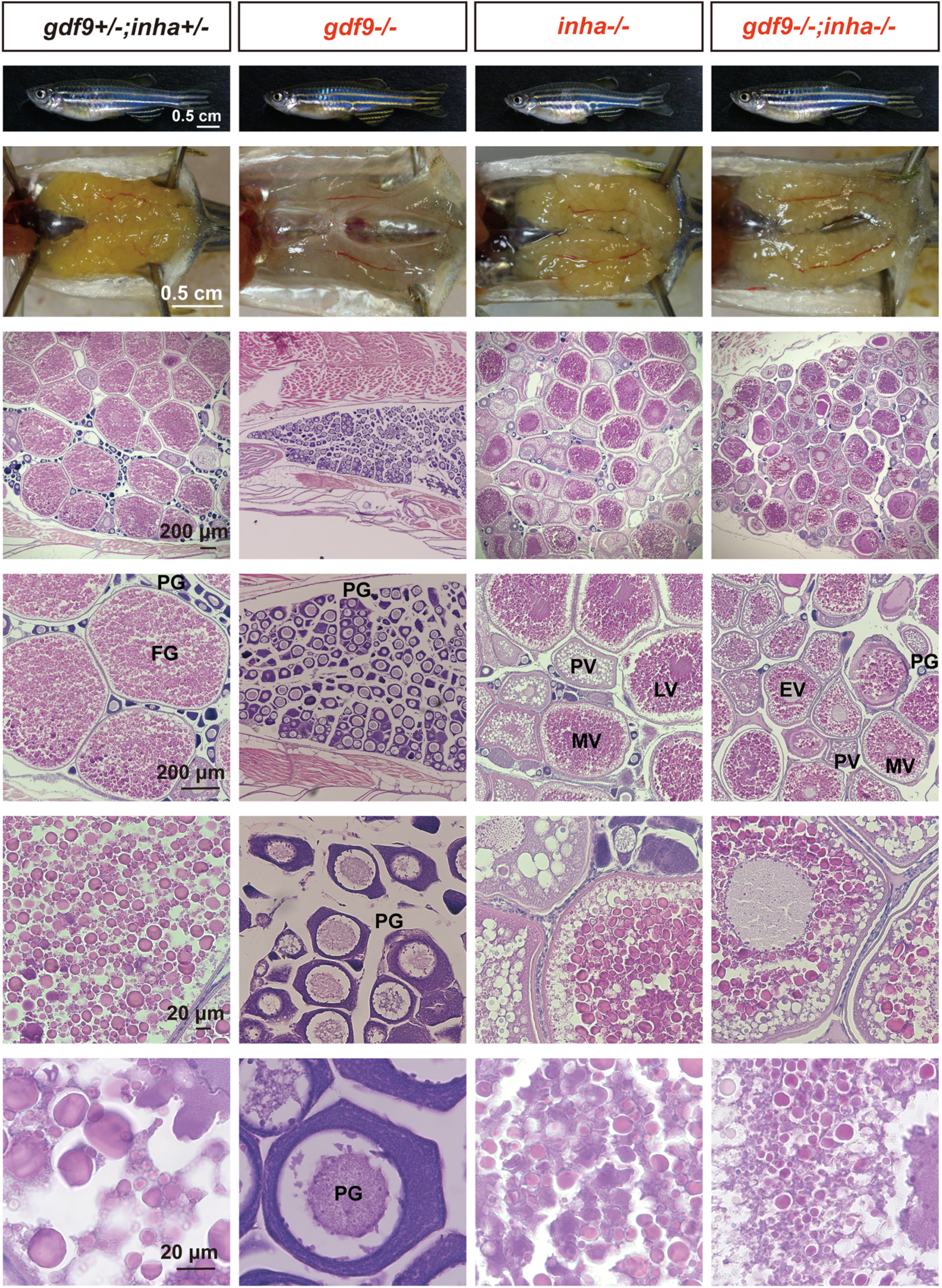
Rescue of follicle blockade in *gdf9* mutant by mutation of inhibin α (*inha*-/-). The follicles in *gdf9* mutant (*gdf9-/-*) were arrested at PG stage at 90 dpf, and double mutation of *gdf9* and *inha* (*gdf9-/-;inha-/-*) resumed follicle activation or PG-PV transition and vitellogenic growth. PG, primary growth; PV, pre-vitellogenic; EV, early vitellogenic; MV, mid-vitellogenic; LV, late vitellogenic; FG, full-grown.

Interestingly, although *inha* mutation could rescue the follicle blockade in the *gdf9* mutant, the restoration of folliculogenesis was not complete. Analysis of follicle composition in the ovaries showed that while folliculogenesis resumed in the double mutant, the follicles could only grow to the mid-vitellogenic (MV) stage (∼450 µm), not the full-grown (FG) stage (>650 µm) as seen in the control and *inha* single mutant (Fig. 8A and B). As the result, the double mutant females were not able to spawn normally with WT males in most fertility tests, similar to the infertility seen in the *inha* mutant (Fig. 8C). When tested in the iSpawner (Techniplast), the double mutant females could sometime release some eggs; however, these eggs were smaller and could not be fertilized (Fig. 8D), consistent with the fact that the follicles could only develop to the MV stage.

**Fig. 8.**
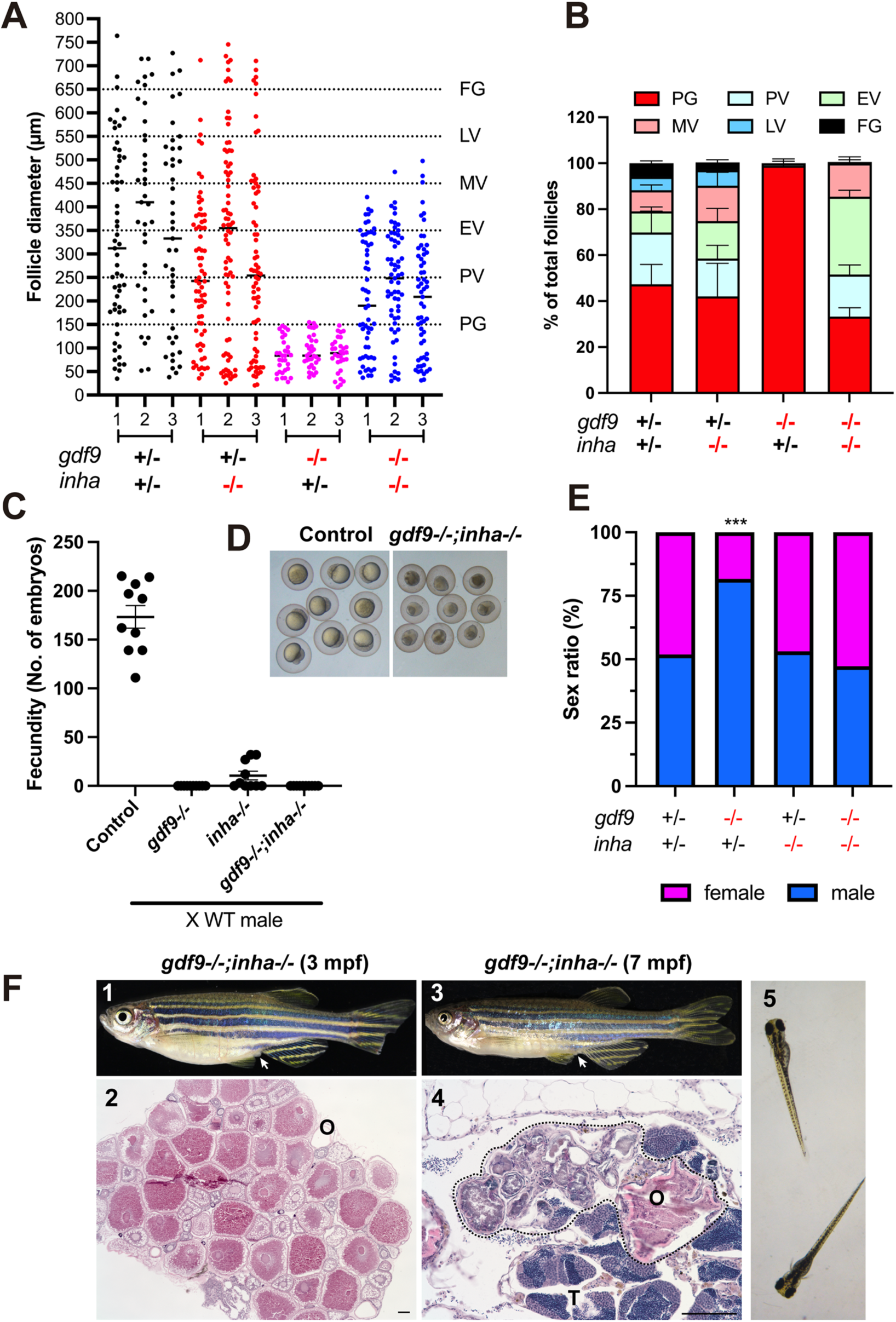
Phenotype analysis of single (*gdf9-/-* and *inha*-/-) and double mutants (*gdf9-/-;inha-/-*) at 90 dpf. (A and B) Follicle composition in different genotypes. The follicles in *gdf9-/-* were arrested at PG stage. Although the PG-PV transition and yolk accumulation resumed in the double mutant (*gdf9-/-;inha-/-*), the vitellogenic growth could not proceed to the FG stage, and the follicles were arrested at the MV-LV transition. (C) Fertility test on females of single (*gdf9-/-* and *inha-/-*) and double mutants (*gdf9-/-;inha-/-*). Five mutant and control females were tested 10 times consecutively on daily basis by natural breeding with WT males (n = 5/test). The number of fertilized eggs or embryos was counted for each female and the data point represents the average of the five fish tested. (D) Eggs spawned by the control (*gdf9+/-;inha+/-*) and double mutant (*gdf9-/-;inha-/-*). The eggs from the double mutant were smaller than those from the control. (E) Prevention of sex reversal in *gdf9-/-* by double mutation with *inha*-/-. The *gdf9-/-* fish showed a male-biased sex ratio (82% males), and this was corrected in the double mutant (47% males). *** P < 0.001 (X^2^ test). (F) Sex reversal of the *gdf9* and *inha* double mutant (*gdf9-/-;inha-/-*) at 7 mpf. (1) Double mutant female at 3 mpf with female-specific genital papilla (arrow); (2) Ovary of 3-mpf double mutant; (3) Sex reversed intersexual fish at 7 mpf with regressed genital papilla (arrow); (4) Ovotestis of 7-mpf double mutant. The follicles in the ovarian tissue (O) were degenerating (circled by dotted line) and the testicular tissue (T) became dominant in the gonads. (5) The offspring from sex reversed double mutant males.

In addition to rescuing follicle blockade, *inha* mutation also delayed sex reversal in the *gdf9* mutant. At 90 dpf, the single mutant (*gdf9-/-*) showed a male-biased sex ratio as described earlier, but this was prevented in the double mutant (*gdf9-/-;inha-/-*), indicating no sex reversal at this time point (Fig. 8E). Despite this, the double mutant females still underwent sex reversal to males at later stage. At 7 months post-fertilization (mpf), most oocytes in the double mutant ovary gradually degenerated and the ovarian tissues were infiltrated by stromal and testicular tissues. The sex reversed males could spawn with WT females to produce normal offspring (Fig. 8F).

### Rescue of inhbaa expression in gdf9 mutant follicles by inha deficiency

The decreased expression of *inhbaa* in *gdf9*-null PG follicles and the rescue of *gdf9* mutant phenotypes by the loss of inhibin (*inha-/-*) both suggested that activin could be involved in Gdf9 actions in controlling early folliculogenesis. To test this hypothesis, we examined the expression of *inhbaa* in PG follicles from *gdf9* single mutant (*gdf9-/-*) and double mutant (*gdf9-/-;inha-/-*), together with *fshr*, *cyp19a1a*, *inhbab* (activin βAb) and *inhbb* (activin βB). As shown in Fig. 9, *fshr*, *cyp19a1a* and *inhbaa* all increased their expression significantly in the PG follicles from *inha-/-* females, in agreement with our recent report [63]. In *gdf9* mutant (*gdf9*-/-), however, *inhbaa* but not *fshr* and *cyp19a1a* showed a significant decrease in expression. Interestingly, in the double mutant (*gdf9-/-;inha-/-*), the expression of all three genes (*fshr*, *cyp19a1a* and *inhbaa*) was raised to the high levels as observed in the *inha* single mutant. The decreased expression of *inhbaa* was therefore not only reversed by the loss of *inha* but increased to a higher level (Fig. 9A-C). By comparison, *inhbab* and *inhbb* did not show any changes in different mutants (Fig. 9D and E). These results suggest a potential role for activin βA, especially *inhbaa,* in mediating actions of oocyte-derived Gdf9.

**Fig. 9.**
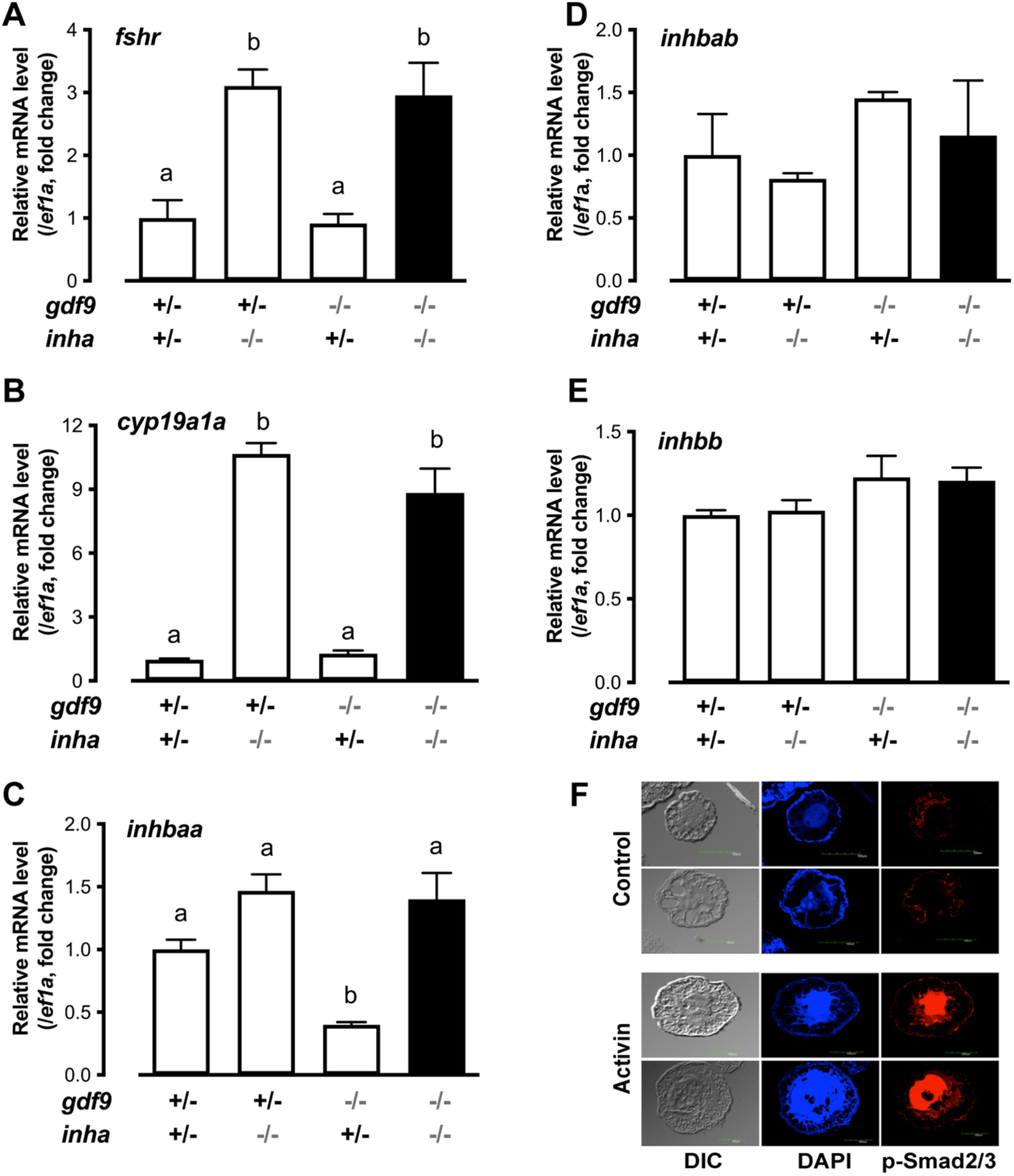
Expression of *fshr*, *cyp19a1a* and activins (*inhbaa*, *inhbab* and *inhbb*) in PG follicles from single (*gdf9-/-* and *inha-/-*) and double mutant (*gdf9-/-;inha-/-*). (A-E) The PG follicles were isolated from the fish of different genotypes at 43 dpf, shortly before puberty onset or PG-PV transition occurs in the control fish (normally around 45 dpf). The expression of *inhbaa* was significantly reduced in *gdf9-/-* mutant; however, it was reversed by *inha* mutation in the double mutant together with *fshr* and *cyp19a1a*, but not *inhbab* and *inhbb*. Different letters indicate statistical significance (P < 0.05). (E) Stimulation of Smad2/3 phosphorylation (p-Smad2/3) in the PV oocytes by activin in vitro. The PV follicles were isolated and treated with recombinant gold activin B (5 U/ml) for 2 h followed by immunofluorescent staining for p-Smad2/3.

Our previous study showed that activin subunits including *inhbaa* were exclusively expressed in the follicle cells whereas activin receptors and its intracellular signaling molecules Smad2/3 were abundantly expressed in the oocyte, suggesting an activin-mediated paracrine pathway in the follicle for follicle cell-to-oocyte signaling [64, 65]. The increased *inhbaa* expression and loss of inhibin in the *gdf9* and *inha* double mutant (*gdf9-/-;inha-/-*) would enhance such signaling by activin. To provide further evidence for direct actions of activin on oocytes, we examined Smad2/3 phosphorylation in PV oocytes in response to activin treatment in vitro. Exposure to activin B (5 U/ml, 2 h) significantly increased the level of p-Smad2/3 in PV oocytes. In addition, the phosphorylated Smad2/3 were mostly concentrated in the germinal vesicles (nuclei), indicating nuclear translocation upon activin stimulation (Fig. 9F).

## Discussion

Folliculogenesis in vertebrates is tightly controlled by both endocrine hormones such as pituitary FSH and LH and local ovarian growth factors [66–68]. In the past decades, evidence has accumulated that local intraovarian paracrine factors play critical roles in orchestrating folliculogenesis [10, 11, 69]. This concept is now being generally accepted largely due to the discovery and characterization of growth differentiation factor 9 (GDF9) [23]. The discovery of GDF9 has triggered tremendous interest in research and numerous studies have demonstrated an essential role for GDF9 in mammalian folliculogenesis [18, 28, 70]. GDF9 is also present in in non-mammalian vertebrates including fish and its oocyte-specific expression seems to be highly conserved [48, 71–77]. However, the functional importance of GDF9 in folliculogenesis remains largely unknown in non-mammalian species.

Using genome editing method TALEN, we successfully knocked out the *gdf9* gene in the zebrafish. Surprisingly, despite its oocyte origin and increased mRNA expression during ovarian differentiation [47], the loss of Gdf9 had no impact on gonadal differentiation as shown by normal sex ratio in young zebrafish in the post-differentiation period. This suggests that although *gdf9* mRNA is expressed early during ovarian formation, its protein may not be translated or secreted until follicle activation at PG-PV transition. In our previous study, we reported that *gdf9* mRNA level was the highest at PG stage and its level declined progressively during zebrafish folliculogenesis. The decline could be the result of increased mRNA turnover and protein translation [48].

Phenotype analysis of zebrafish *gdf9* mutant (*gdf9*-/-) showed that similar to GDF9 in the mouse, Gdf9 also plays an essential role in controlling early folliculogenesis in fish. The disruption of *gdf9* gene caused a complete arrest of follicle development at the PG stage without any signs of follicle activation to enter the fast-growing SG phase. The follicles could grow to nearly full size of PG stage; however, further development to PV stage, which is the beginning of the SG phase characterized with the formation of cortical alveoli in the oocytes, was completely blocked. Since the appearance of cortical alveoli is considered a marker for follicle activation and puberty onset [55], the arrest at PG-PV transition also resulted in failed puberty onset and female maturation, and therefore infertility. Our observation in zebrafish agrees well with the report in the mouse model. In *Gdf9*-null female mice, the primordial and one-layer primary follicles could form normally; however, further development to two-and multiple-layer secondary follicles was blocked [28]. The blockade at the primary-secondary follicle transition in *Gdf9-*null mice is surprisingly similar to the blockade at the PG-PV (primary-secondary growth) transition in zebrafish *gdf9* mutant. Another striking similarity between *Gdf9*-null mice and *gdf9*-null zebrafish is the lack of cortical alveoli (granules) in the oocytes [28]. These results strongly suggest that the fundamental functions of GDF9 in controlling early follicle development are high conserved in vertebrates. Our results, however, contrast with a recent study in zebrafish, which reported no phenotypes in *gdf9*-null zebrafish [78]. The reason for such discrepancy is unknown. Similar situations have also been reported in mice. For example, the *Sirt1*-null mice generated by different groups exhibited different phenotypes, which might be due to different genetic backgrounds of the mice used [79, 80].

In addition to controlling PG-PV transition or follicle activation, our data also suggest an important role for Gdf9 in maintaining feminization including the ovary although it was not involved in primary gonadal differentiation. The loss of Gdf9 caused gradual degeneration of the follicles followed by transformation of the ovary to testis, resulting in all-male phenotype in the end. This process seems due to the change of dominance of the two antagonistic sex differentiation pathways, which are present in each individual of vertebrate species [81]. The loss of Gdf9 weakens the female pathway in zebrafish, resulting in a switch of dominant pathway from female to male. This idea is supported by our evidence that deletion of *dmrt1*, a critical gene in male-promoting pathway [82], prevented sex reversal in the *gdf9* mutant. Interestingly, although *gdf9* expression could also be detected in adult testis [48], its loss had no impact on spermatogenesis in males. In *Gdf9*-null mice, the oocytes arrested at the primary follicle stage also underwent degeneration followed by collapse of the zona pellucida and luteinization of the granulosa cells [28], but not sex reversal as we observed in zebrafish.

Our observation on *gdf9*-null zebrafish suggests that although pituitary gonadotropins play pivotal roles in driving follicle growth and maturation [1, 2], an intrinsic regulatory mechanism exists in the follicle that orchestrates folliculogenesis in zebrafish, and that oocytes play an active role in timing the developmental process. As an oocyte-derived factor, Gdf9 is expected to act on the surrounding follicle cells where it may interact with the endocrine hormones and other local factors to control follicle development. The exact functions of Gdf9 in zebrafish follicles are largely unknown. Future studies using recombinant zebrafish Gdf9 and in vitro follicle or follicle cell culture will shed light on the biological activities of Gdf9 in the follicle. Using this approach, we recently demonstrated that recombinant zebrafish Gdf9 could activate the Smad2 signaling pathway in cultured follicle cells and it significantly increased the expression of activin subunits (*inhbaa* and *inhbb*) but decreased that of anti-Müllerian hormone (*amh*), a male-promoting factor in the gonads [47]. It is therefore conceivable that the oocyte-derived Gdf9 may act on the follicle cells to stimulate the factors that promote follicle growth and maintenance such as activin but suppress those involved in masculinization such as Amh, which increases its expression significantly during female-to-male sex reversal [2].

Follicle activation or PG-PV transition is a critical stage in zebrafish folliculogenesis, which involves multiple endocrine, paracrine and autocrine factors as well as various signaling pathways [62]. Pituitary gonadotropins (FSH and LH) are undoubtedly the master hormones that control this process. The FSH receptor (*fshr*), which is co-activated by both FSH and LH in zebrafish [83], increases its expression significantly during PG-PV transition [58, 84], suggesting an important role for gonadotropin signaling in this event. This has recently been confirmed by the observation that deletion of pituitary *fshb* gene (FSHβ) significantly delayed the PG-PV transition and therefore puberty onset [1] and knockout of *fshr* suppressed ovarian growth and completely blocked follicle activation [2, 85]. Like those in mammals, local ovarian factors are also implicated in controlling follicle activation in zebrafish. Activin subunits especially *inhbaa* increased expression significantly during the PG-PV transition [61, 62]. Similarly, epidermal growth factor (EGF) family ligands including EGF (*egf*) and transforming growth factor α (TGF-α/*tgfa*) as well as their receptor EGFR (*egfra*) also showed an increased expression at the PG-PV transition [64]. In zebrafish follicles, EGF family ligands are primarily expressed in the oocyte whereas their receptor *egfra* is mainly expressed in the somatic follicle cells [64], which respond to EGF strongly by MAPK phosphorylation, increased expression of activin subunits (*inhbaa*, *inhbab* and *inhbb*), and decreased expression of follistatin (an activin binding protein) [86, 87]. Like Gdf9, EGF ligands and EGFR may represent another paracrine pathway within the follicle that mediates signaling from oocyte to follicle cells. Our recent study showed that disruption of *egfra* in zebrafish also blocked follicle development at PG-PV transition followed by sex reversal to males [59], which is similar to the phenotypes of *gdf9* mutant observed in the present study. These results indicate that follicle development in zebrafish ovary is finely controlled by multiple factors from the oocyte and that the regulation of folliculogenesis involves many checkpoints for regulation by both internal and external signals. This view is further supported by a recent study in zebrafish showing that mutation of BMP15 (*bmp15*), also an oocyte-specific factor from TGF-β family, caused an arrest of follicle growth at PV stage [78], in contrast to the arrest at PG stage as seen in the *gdf9* and *egfra* mutants.

Having defined the functional role and importance of Gdf9 in controlling folliculogenesis in zebrafish, we went on to explore the mechanism underlying its actions. Much needs to be learnt to unravel how GDF9 acts, which remains an issue in mammalian models as well [88]. One important clue from our experiments was the dramatic decrease in the expression of activin subunit *inhbaa* in *gdf9* mutant follicles. This agrees well with our previous in vitro study showing stimulation of *inhbaa* expression in cultured follicle cells by recombinant zebrafish Gdf9 [47]. This has led us to hypothesize that part of the mechanism for Gdf9 to regulate follicle development is to increase the production of activin in the follicle cells, which in turn acts on the oocyte that expresses activin receptors and signaling molecules Smad2/3/4 [65]. Our observation that the PV oocytes responded strongly to activin treatment in terms of Smad2/3 phosphorylation and nuclear translocation further supports the idea that the oocyte is a direct target for activin actions. The loss of oocyte-derived Gdf9 would lead to decreased expression of activin subunits, therefore compromising an important signaling from the follicle cells to oocyte and therefore resulting in the arrest of follicle development.

As a key local factor from the follicle cells targeting the oocyte, activin has been proposed to play a central role in mediating both endocrine hormones such as gonadotropins [89] and local paracrine factors such as EGF ligands [86, 87] in the follicle. Similarly, activin may also play a role in mediating Gdf9 actions. In support of this hypothesis was our evidence that knockout of *inha* could partially reverse the phenotypes of *gdf9* mutant. Inha is also a member of TGF-β family and it dimerizes with an activin β subunit (αβ) to form inhibin, which acts as an activin antagonist. The loss of *inha* advanced PG-PV transition and therefore puberty onset in female zebrafish [63], in contrast to the PG-PV blockade in *gdf9* mutant. Double mutations of *inha* and *gdf9* (*gdf9-/-;inha-/-*) not only resumed follicle growth to MV stage with cortical alveoli formation and yolk accumulation, but also delayed sex reversal of *gdf9-/-* females to males. Our discovery is surprisingly similar to a previous study in mice. The loss of *Gdf9* in mice blocked follicle development at the primary follicle stage without thecal layers [28]. Interestingly, double mutant of *Gdf9* and *Inha* partially rescued the phenotypes of *Gdf9* single mutant. The follicles in the double mutant ovary (*Gdf9-/-;Inha-/-*) contained not only primordial and one-layer primary follicles, but also two- and multiple-layer secondary follicles with thecal layers [90]. Interestingly, although EGF also stimulated expression of all activin subunits in zebrafish follicle cells [87] and the loss of *egfra* gene led to similar phenotypes to those of *gdf9* mutant, viz. blockade at PG-PV transition followed by sex reversal to males, *inha* mutation could not rescue the phenotypes of *egfra* mutant [59], suggesting differential mechanisms underlying actions of Gdf9 and EGF family ligands.

How the loss of inhibin rescued the defective phenotypes of *gdf9* mutant is not clear. Interestingly, three genes (*fshr*, *cyp19a1a* and *inhbaa*) that are considered functionally important in promoting folliculogenesis were all increased dramatically in the double mutant (*gdf9-/-;inha-/-*) to the high levels seen in *inha* single mutant. The increased expression of these genes would suggest increased gonadotropin signaling, estrogen biosynthesis and activin production, all of which could enhance follicle development, therefore explaining how mutation of *inha* could rescue the defects of *gdf9* mutant. Activin is the most interesting among the three in that *inhbaa* was the only gene that showed a significantly decreased expression in *gdf9*-null PG follicles and its expression rebounded dramatically in the double mutant (*gdf9-/-;inha-/-*). In addition to increased expression, the bioactivity of activin is also expected to increase in the *inha* mutant for two reasons. First, as a potent activin antagonist, the loss of inhibin is expected to cause a reduced antagonism against activin. Second, sharing the same β subunits, activin and inhibin compete for β subunits in their biosynthesis; therefore, the loss of inhibin α subunit (*inha*) is expected to increase the formation of activin molecules. Whether the rescue of *gdf9* mutant phenotypes by *inha* mutation involves activin alone or its interaction with gonadotropins and estrogens would be an interesting issue to explore in future studies. As for *cyp19a1a*, our recent study has provided clear evidence for its critical role in ovarian differentiation; however, *cyp19a1a* was not involved in promoting PG-PV transition [57], which agrees with the finding in medaka fish [91]. This view is further supported by our evidence that treatment with E2 could not overcome the blockade of follicle development in the *gdf9* mutant although E2 could prevent sex reversal and induce vitellogenin production in the liver as expected.

In summary, using genome editing method, we performed a genetic analysis on roles and functional importance of *gdf9* in zebrafish folliculogenesis. Our data demonstrated that *gdf9* is critical for early follicle development, especially at follicle activation or transition from PG to SG phase. The loss of *gdf9* resulted in a complete cessation of follicle development at the PG stage. Gene expression analysis especially genetic analysis with *inha* and *gdf9* double mutant provided critical insight into how Gdf9 works in zebrafish follicles. We hypothesize that the oocyte-derived Gdf9 works on the surrounding follicle cells to stimulate biosynthesis of activins, which in turn act back on the oocyte in a paracrine manner to activate the Smad2/3 signaling pathway, promoting formation of cortical alveoli and therefore follicle activation (Fig. 10). This study provides a strong support to the view that oocytes play active and important roles in orchestrating folliculogenesis and Gdf9 has conserved functions across vertebrates.

**Fig. 10.**
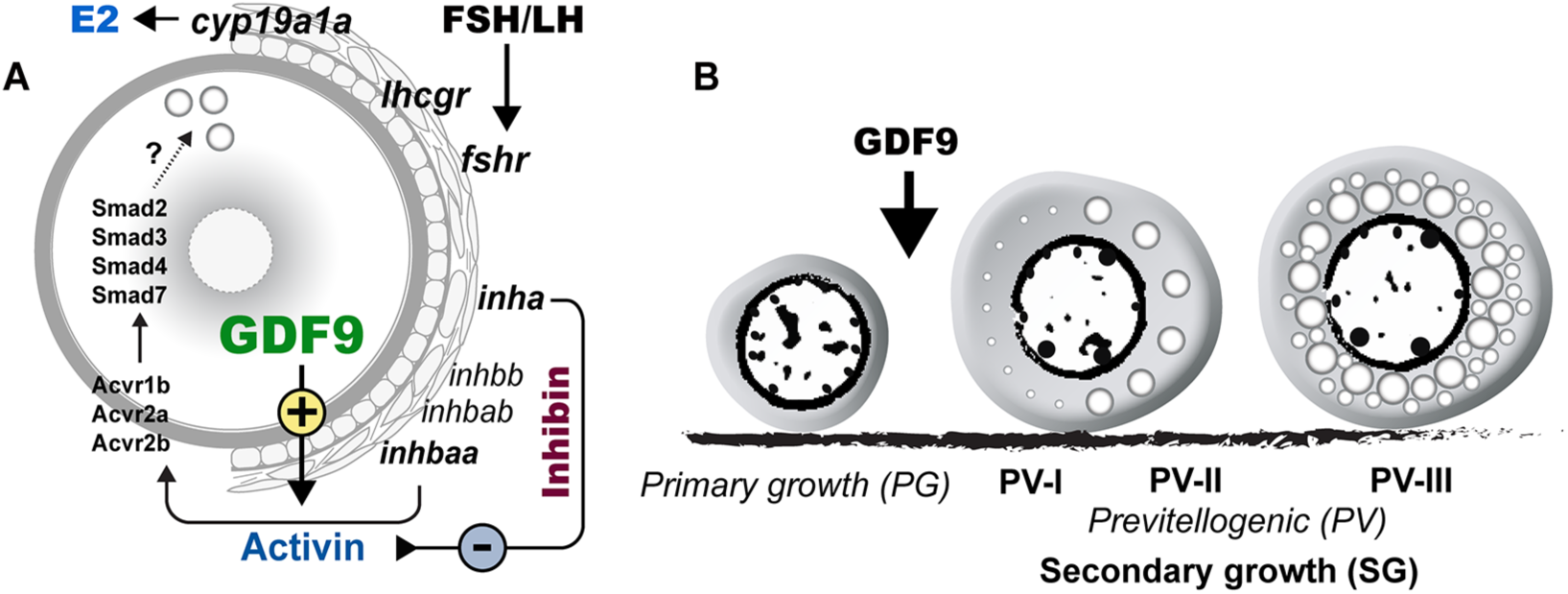
Hypothetical model on roles of oocyte-derived GDF9 (*gdf9*) in controlling zebrafish folliculogenesis. (A) Intrafollicular distribution of GDF9 and other important genes (*fshr*, *lhcgr*, *cyp19a1a*, *inhbaa*/*ab*, *inhbb*, *inha*, *acvr1b*, *acvr2a*/*b* and *smad2/3/3/7*). (B) GDF9 from the oocyte plays an important gating role in controlling zebrafish follicle activation or PG-PV transition and therefore puberty onset. PV-I, early PV with one single layer of small cortical alveoli; PV-II, mid-PV with one single layer of large cortical alveoli; PV-III, late PV with multiple layers of cortical alveoli.

## Materials and Methods

### Animals

The AB strain of wild type (WT) zebrafish (*Danio rerio*) was used to generate mutant lines, and the fish were kept in the flow-through ZebTEC multilinking zebrafish system (Tecniplast, Buguggiate, Italy) on 14L:10D lighting cycle. The system condition was maintained as follows: temperature 28°C, pH 7.3 and conductivity 400 mS/cm. The fish were fed twice per day with Otohime fish diet (Marubeni Nisshin Feed, Tokyo, Japan), which was delivered by the Tritone automatic feeding system (Tecniplast). All experiments were endorsed by the Research Ethics Panel of University of Macau.

### Generation of gdf9 mutant zebrafish

To create mutant zebrafish, the whole sequence containing *gdf9* gene (ID: ENSDARG00000003229) was retrieved from the Ensembl database for target site identification. The sequence in the first exon downstream of the ATG start codon was chosen for targeting by TALEN. Both left and right TALEN arms were designed by the online software (TAL Effector Nucleotide Targeter 2.0 Tools; https://tale-nt.cac.cornell.edu/node/add/talen) as we previously reported [1], and the sequences are: TAGTGCGCTTTGTTACC (left TALE), TTGTCCTCGCAGTTCTCGGA (right TALE), and TACCCAGATCATTAA (spacer sequence) (Fig.1). Gene-specific TALEN constructs were assembled using the TALEN Golden Gate assembly system as described [92] with the two backbone plasmids pCS2TAL3DD and pCS2TAL3RR (Addgene, Cambridge, MA). The TALE mRNAs were generated by in vitro transcription using the mMESSAGE mMACHINE SP6 kit (Ambion, Austin, TX), and co-injected (50 pg each) into one or two-cell stage embryos. To detect mutations, the genomic DNA was extracted from each embryo or caudal fin cut and analyzed by high resolution melt analysis (HRMA) as we previously reported [12]. The primers for HRMA analysis are listed in Table 1. The reaction was performed with EvaGreen Supermix on the CFX96 real-time PCR machine, and the data were analyzed with the precision melt analysis software (Bio-Rad, Hercules, CA). The mutations were confirmed by sequencing (Tech Dragon, Hong Kong).

**Table 1.**
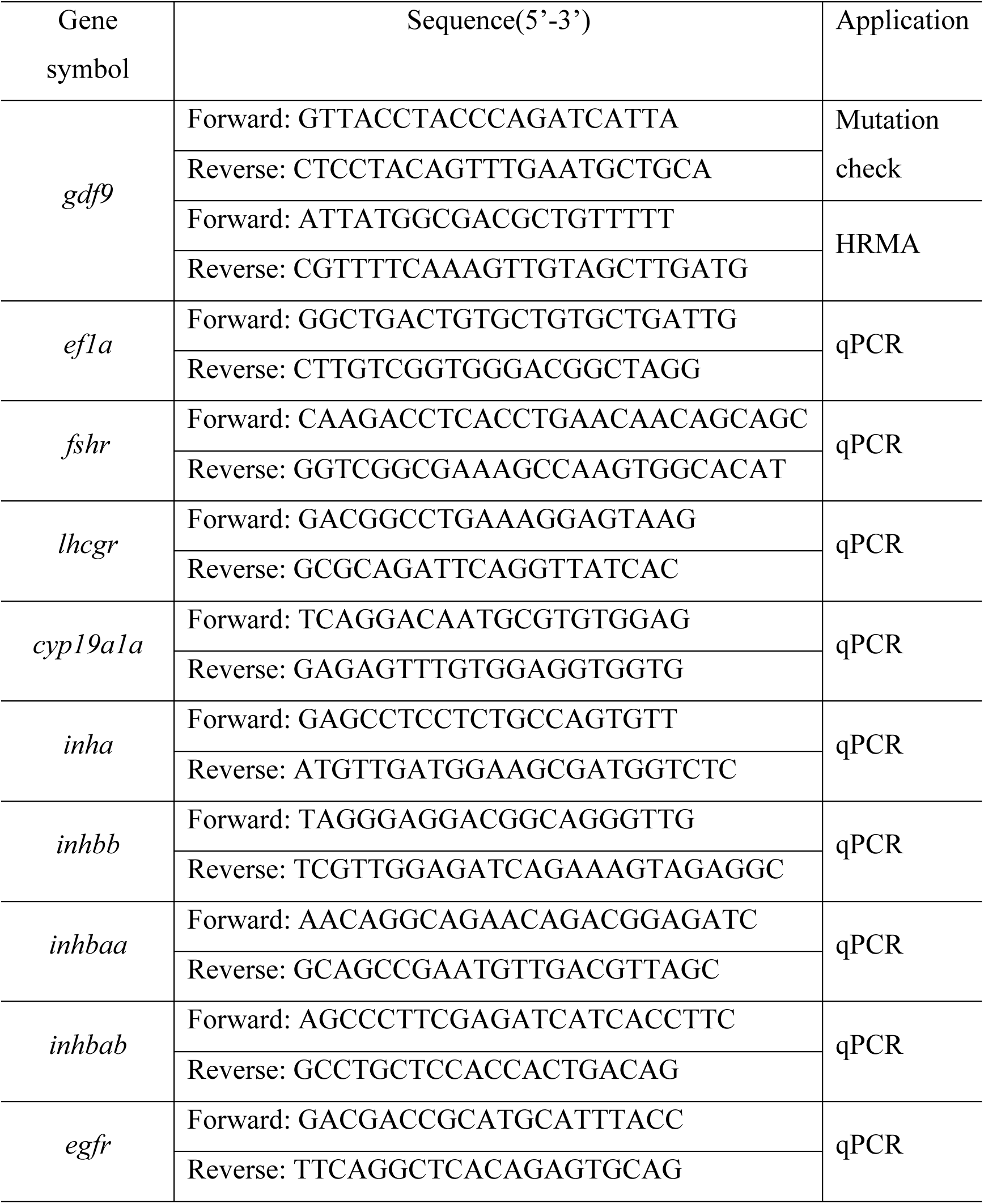
Primer list

### Genomic DNA isolation

Genomic DNA was isolated from embryos or caudal fins of WT or mutant fish as described [1]. Briefly, a piece of caudal fin or an embryo was incubated in 40 μl NaOH (50 mM) at 95℃ for 10 min. After cooling down to room temperature, 4 μl Tris-HCl (pH8.0) was added to each sample. After centrifugation, the supernatant was then used directly as the template for HRMA.

### Real-time qPCR quantification

Total RNA was isolated from the follicles with Tri-Reagent (Molecular Research Center, Cincinnati, OH) according to the protocol of the manufacturer and our previous reports [48, 93]. The RNA was reverse transcribed into cDNA at 42°C for 2 h in a total volume of 10 μl reaction solution that contains total RNA, reverse transcription buffer, DTT, oligo-dT and reverse transcriptase (Invitrogen, Hercules, CA).

The expression levels of target genes in the follicles from WT or mutant fish were determined by real-time qRCR as described [93]. The standard for each gene was prepared by PCR amplification of cDNA fragments with specific primers (Table 1). These amplified amplicons were used to construct standard curves in the assay. Real-time qPCR was carried out on the CFX96 Real-Time PCR Detection System (Bio-Rad) in a volume of 20 μl that contained 10 μl diluted RT reaction mix, 1× PCR buffer, 0.2 mM of each dNTP, 2.5 mM MgCl_2_, 0.2 μM of each primer, 0.75 U Taq polymerase, 0.5× EvaGreen (Biotium, Hayward, CA). The reaction profile consisted of 38 cycles of 94°C for 30 sec, 60°C for 30 sec, 72°C for 1 min, and 80°C for 7 sec for signal detection. A melt curve analysis was performed at the end of amplification to demonstrate reaction specificity.

### Histological analysis of gonads

The gonadal development and morphology were examined by histology as described [55]. Briefly, the body trunk of each fish was fixed in Bouin’s solution overnight at room temperature. Following dehydration and embedding in paraffin, the samples were sectioned on a Leica microtome (Wetzlar, Germany) and stained with hematoxylin and eosin (H&E). For the convenience of analysis, we divide zebrafish follicle development into two phases: primary growth (PG) (<150 µm; stage I) and secondary growth (SG) (∼150-750 µm), and the SG phase is further divided into pre-vitellogenic (PV, ∼250 µm; stage II), early vitellogenic (EV, ∼350 µm; early stage III), mid-vitellogenic (MV, ∼450 µm; mid-stage III), late vitellogenic (LV, ∼550 µm; late stage III) and full-grown (FG, ∼650 µm; full-grown stage III) stages, based on size and morphological markers such as cortical alveoli and yolk granules [84, 94]. To quantify follicle composition in the ovary, we performed serial longitudinal sectioning of the whole fish at 5 µm and measured diameters of follicles on three largest sections spaced at least 60 µm apart by the NIS-Elements BR software (Nikon, Tokyo, Japan). To ensure the accuracy of diameter measurement for follicle staging, we only measured the follicles with visible nuclei (germinal vesicles) on the section.

### Estradiol treatment in vivo

To examine if the phenotype of *gdf9* mutant could be rescued by estrogens, we treated the mutant fish with E2 (Sigma-Aldrich, St. Louis, MO) followed by sampling for histological analysis. Two approaches were used for estrogen treatment: water-borne exposure and oral administration by natural feeding. For water-borne exposure, E2 stock in ethanol was added to the water in fish tank to the final concentration of 10 nM, and the juvenile fish were treated for 20 days from 40 to 60 dpf. The water was replaced by half every day with E2 supplement with ethanol being used as the vehicle control. For oral administration, the mutant females were treated with E2 for 17 days from 45 to 62 dpf. In brief, the fish were fed twice a day with E2-containing Otohime fish diet (0 or 2 μg g-1), each at 5% of total fish body weight in the tank (10% per day in total), with supplementation of brine shrimp larvae. During the treatment period, the water was renewed daily to maintain good water quality.

### SDS-PAGE examination for vitellogenin production

The whole blood was collected from each fish by cutting the tail. The blood samples were left at room temperature for clotting followed by centrifugation to obtain the serum. The sera were subject to standard sodium dodecyl sulfate (SDS) PAGE analysis (1 µl per lane) for vitellogenin proteins. We also examined vitellogenin production in the liver. Briefly, the liver was dissected from each fish and lysed by in 400 µl SDS sample buffer (62.5 mM Tris-HCl [pH 6.8] at 25°C, 1% [wt/vol] SDS, 10% glycerol, 5% 2-mercaptoethanol). All samples were heated to ∼95°C for 10 min, cooled on ice, centrifuged for 5 min, and then loaded to 12% SDS-PAGE gel for electrophoresis.

### Immunofluorescent staining for Smad2/3 phosphorylation

To demonstrate responsiveness of oocytes to activin, PV follicles were isolated from the ovary and incubated in 60% Leibovitz L-15 medium (Gibco, Thermo Fisher Scientific, Waltham, MA) in the presence or absence of recombinant goldfish activin B (5 U/ml, 2 h) prepared in our laboratory as reported [95, 96]. The follicles were fixed after incubation in 2% formaldehyde–phosphate-buffered saline (PBS), permeabilized with 0.02% TritonX-100 in PBS, incubated with primary anti-p-Smad2/3 (Cell Signaling, Danvers, MA) and then secondary goat anti-rabbit IgG conjugated with Alexa Fluor 594 (Thermo Fisher Scientific). The follicles were mounted on a slide for observation with the FluoView FV1000 IX81 confocal microscope (Olympus, Tokyo, Japan).

### Data analysis

The expression levels of target genes were normalized to the house keeping gene *ef1a*. All values were expressed as the mean±SEM, and the data were analyzed by Dunnett’s Multiple Comparison Test with Prism (GraphPad Software, San Diego, CA).

## Acknowledgements

We thank Ms. Phoenix Un Ian LEI for the maintenance and management of the zebrafish facility and the Histology Core of the Faculty of Health Sciences for technical support. This study was supported by grants from the University of Macau (MYRG2017-00157-FHS, MYRG2019-00123-FHS and CPG2020-00005-FHS) and The Macau Fund for Development of Science and Technology (FDCT173/2017/A3 and FDCT0132/2019/A3) to W.G. KW is supported by the Macau Young Scholars Program (AM2020025).

## Notes

### Competing Interest Statement

The authors have declared no competing interest.

